# Tubulin Acetylation Promotes Penetrative Capacity of Cells Undergoing Radial Intercalation

**DOI:** 10.1101/2021.04.29.441997

**Authors:** Caitlin Collins, Sun K. Kim, Rosa Ventrella, Jen W. Mitchell, Brian Mitchell

## Abstract

The post-translational modification of tubulin provides a wide diversity of differential functions to microtubule networks. Here we address the role of tubulin acetylation on the penetrative capacity of cells undergoing radial intercalation in the skin of *Xenopus* embryos. Radial intercalation is the process by which cells move apically and penetrate the epithelial barrier via inserting into the outer epithelium. As such there are two opposing forces that regulate the ability of cells to intercalate: the restrictive forces of the epithelial barrier versus the penetrative forces of the intercalating cell. By positively and negatively modulating tubulin acetylation specifically in the intercalating cells, the timing of intercalation can be altered such that cells with more acetylated microtubules penetrate the epithelium faster. Moreover, the *Xenopus* epithelium is a complex array of variable types of vertices and we find that intercalating cells preferentially penetrate at higher order “rosette” vertices as opposed to the more prevalent tricellular vertices. We observed differential timing in the ability of cells to penetrate different types of vertices, indicating lower order vertices represent more restrictive sites of insertion. Interestingly, we are able to shift the accessibility of early intercalating cells towards the more restrictive tricellular junctions by modulating the level of tubulin acetylation and the subsequent penetrative capacity of intercalating cells. Overall our data implicate tubulin acetylation in driving tissue penetration of intercalating cells.

## Introduction

Epithelia represent a barrier between different tissues or the external environment. Critical to this function they must maintain epithelial integrity during a diverse array of insults including injury and disease. In contrast, the ability of cells to penetrate epithelial barriers is important for proper development and tissue homeostasis. The balance between maintaining epithelial integrity and allowing cell penetration is a complex interplay involving cell signaling, junctional remodeling, directed cell movements, and cytoskeletal dynamics. Here we address the role of tubulin acetylation on both the timing and the vertex accessibility of cells undergoing radial intercalation into the epithelium of *Xenopus* embryonic skin.

During *Xenopus* development, both multiciliated cells (MCCs) and ionocytes (ICs) differentiate in a sublayer of the epithelium. In order to become functional, these cells must undergo a short apical migration and insert into the outer epithelium in a process called radial intercalation. Radial intercalation involves multiple steps requiring cells to 1.) migrate in the apical direction, 2.) penetrate the epithelial barrier by apically inserting between outer epithelial cells, and 3.) expanding their apical surface until they achieve proper size (Collins et al., 2020b). Cytoskeletal forces are known to be integral cell autonomous features of the intercalation process. Actin-based forces, as well as small small GTPase activity, drive apical expansion of MCCs (Ioannou et al., 2013; Kim et al., 2012; Kulkarni et al., 2018; Sedzinski et al., 2016, 2017). Additionally, we have previously shown that centriole number and the subsequent changes to microtubule (MT) accumulation can regulate the timing of apical insertion, such that cells with more centrioles or more MTs insert earlier than cells with less (Collins et al., 2020a). While these results clearly establish a critical role for MTs during intercalation, the functional mechanism of how MTs promote apical insertion remains poorly understood.

MTs have numerous cellular functions, including providing the tracks for intracellular motor-based trafficking, facilitating the generation of cellular polarity, and providing the physical forces for diverse processes (e.g. mitosis). It is well established that MTs play critical roles during cellular migration including facilitating the transport of membrane bound vesicles and signaling molecules to the leading edge (Etienne-Manneville, 2013). Most MT-based functions require MT directionality. We have previously reported that MTs are essential for regulating the timing of apical insertion during intercalation but importantly an increase in MTs in any orientation promotes insertion (Collins et al., 2020a). These results suggest that MTs may have a critical function independent of vesicle trafficking. Tubulin acetylation is known to promote the stabilization of MTs and lead to longer lived MTs which are more resistant to strain and more easily repaired (Janke and Montagnac, 2017; Xu et al., 2017). Additionally, acetylated MTs promote motor based trafficking (Alper et al., 2014; Nekooki-Machida and Hagiwara, 2020; Reed et al., 2006). Furthermore, the acetylation of MTs has known roles in promoting cell migration and invasive behavior in numerous contexts, most notably in collective cell migration where it promotes focal adhesion stability (Bance et al., 2019; Boggs et al., 2015; Castro-Castro et al., 2012). Here we test the hypothesis that acetylated MTs modulate both the timing of apical insertion and the penetration through more restrictive lower order vertices.

Radial intercalation has been primarily addressed from the perspective of the intercalating cell, yet the surrounding tissue environment likely contributes significantly to the process. Long range cell migration events require chemo-attractant and chemo-repulsive signals (Szabo and Mayor, 2018). In fact, during *Xenopus* epiboly when cells radially intercalate to promote tissue thinning, cells respond to the chemo-attractant C3a (Szabo et al., 2016). However, there is no evidence to date that the short single cell layer radial intercalation of MCCs and ICs into the skin requires external cues. Furthermore, tissue features such as substrate stiffness are known to contribute to collective cell movements in multiple contexts, including neural crest migration, but are unlikely to contribute significantly to the short individual cell movements of radial intercalation (Barriga et al., 2018; Barriga and Mayor, 2019; Szabo and Mayor, 2018; Zanotelli et al., 2019). Additionally, cellular arrangements provide a variety of tissue topologies that could offer differential access to intercalating cells. In the drosophila oocyte it has recently been shown that migrating border cells prefer a central path through the egg chamber that represents the topographical path of least resistance via increased spacing between cells (Dai et al., 2020). In *Xenopus* skin it has been reported that intercalating cells penetrate exclusively at multi-cellular vertices rather than between two cells (Stubbs et al., 2006). This implies that vertices represent weak spots in the epithelium. Moreover, it is known that during the massive tissue remodeling that occurs during convergent-extension elongations, that the sites of active movement are often associated with rosette vertices, sites where 5 or 6 cells come together (Blankenship et al., 2006; Lienkamp et al., 2012; Vanderleest et al., 2018). This predilection for higher order vertices suggests that these represent more pliable sites of epithelial transformation. However, how the variability of vertex number affects the ability of cells to penetrate during radial intercalation remains unexplored. Here we address the preference of MCCs for intercalating between higher order vertices (e.g. rosettes) and the malleability of that preference towards the more restrictive lower order tricellular vertices based on modulating the penetrative capacity of the intercalating cell.

## Results and Discussion

MTs are important for the intercalation of both MCCs and ICs. Increasing centriole number results in an increase in MTs and precocious apical insertion (Collins et al., 2020a). In contrast, treatment with the MT depolymerizing drug Nocodazole leads to partial intercalation where cells do not achieve normal apical size (Werner et al., 2014). Here we have built on these results by performing an analysis of the timing of apical insertion in the presence of Nocodazole. Radial intercalation is a progressive process but we have previously defined the initial apical insertion event as the point when cells achieve a small apical size of 35μm^2^, equivalent to the average size of the apical domain at the earliest stage observed (Collins et al., 2020a). Here we show that cells treated with Nocodazole are delayed specifically in apical insertion such that at each developmental stage (ST) significantly fewer MCCs or ICs had successfully penetrated the epithelial barrier (Supplemental Figure 1). Surprisingly, we have observed that the majority of MTs found in intercalating cells are acetylated (Werner et al., 2014). Acetylated MTs have been reported to be more stable and more resistant to strain leading us to hypothesize that the acetylation of MTs provides added strength to the MT network that facilitates their ability to penetrate through the epithelial barrier during apical insertion.

To test our hypothesis, we first overexpressed (OE) the deacetylase HDAC6 specifically in post-mitotic MCCs using expression via the alpha tubulin promoter (Tub) (Stubbs et al., 2006). We did not see a significant difference in the overall MT network as quantified by fluorescent intensity of antibody staining using a beta-tubulin antibody (Figure 1A, C). In contrast, we did observe a significant decrease in tubulin acetylation by acetylated tubulin antibody staining in MCCs overexpressing HDAC6 (Figure 1B, D). Importantly, when we performed a time course of MCC intercalation we found that there was a significant delay in apical insertion with fewer cells having successfully penetrated the outer epithelium at each time point (Figure 1E, F). Ultimately the typical number of MCCs intercalate, but the delay in apical insertion suggests that tubulin acetylation and the corresponding change in MT rigidity is important for providing the protrusive ability required for MCCs to quickly penetrate the outer epithelium.

**Figure 1.**
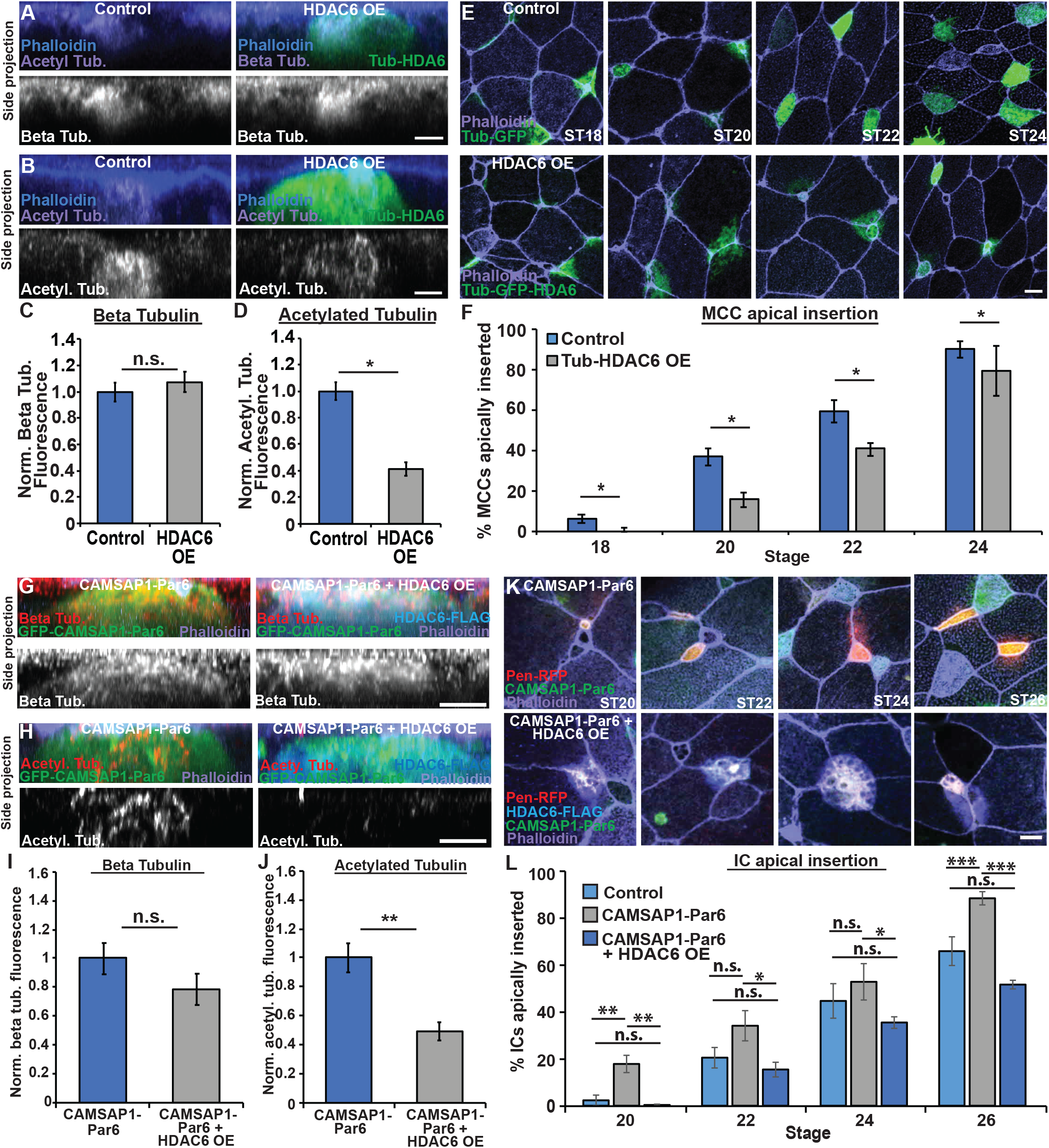
Loss of MT acetylation delays apical insertion. **A-B,** Side projections of intercalating control and Tub-HDAC6 OE MCCs fixed and stained with phalloidin and α-beta tub **(A)** or α-acetyl. tub. **(B)**. **C**-**D,** Quantification of beta tub **(C)** and of acetyl. tub. **(D)** in control and HDAC6 OE MCCs. Fluorescence was normalized relative to control (uninjected) MCCs in mosaic embryos for each experiment. **E,** Z-projections of embryos mosaically injected with Tub-GFP or Tub-GFP-HDAC6 DNA and phalloidin to assay apical insertion. **F,** Quantification of the percentage of MCCs apically inserted at each stage. **G-H,** Side projections of intercalating CAMSAP1-Par6 and CAMSAP1-Par6 + HDAC6 OE ICs stained with α-FLAG, phalloidin, and α-beta tub. **(G)** or α-acetyl. tub. **(H)**. **I-J,** Quantification of beta tub. **(I)** or acetyl tub. **(J)** in CAMSAP1-Par6 and CAMSAP1-Par6 + HDAC6 OE ICs. Fluorescence was normalized relative to (control) ICs expressing only CAMSAP1-Par6 in mosaic embryos for each experiment. **K,** Z-projections of embryos mosaically injected with Pen-RFP and HDAC6-FLAG DNA and CAMSAP1-Par6 mRNA and stained with α-FLAG and phalloidin to assay apical insertion. **L,** Quantification of the percentage of ICs apically inserted at each stage. For all bar graphs, bars represent the average, error bars indicate SD, and *p<0.05, **p<0.01, ***p<0.001. Analysis includes n > 80 cells from at least 6 embryos per condition (C), n > 60 cells from at least 5 embryos per condition (D) and n > 175 cells from at least 9 embryos per condition/time point (F), n > 15 cells from at least 5 embryos per condition (I), n > 25 cells from at least 7 embryos per condition (J), n > 50 cells from at least 9 embryos per condition/time point (L). Scale bars in A, B, G and H is 5μm and in E and K is 10μm.

ICs have considerably fewer MTs than MCCs and are slower in their progression through radial intercalation, with the primary delay being in apical insertion. We have shown that expression of the MT (-) end protein CAMSAP1 fused to the apically positioned Par6 protein (CAMSAP1-Par6) increases MTs in ICs and is sufficient to drive precocious apical insertion (Collins et al., 2020a). Here we have addressed whether the acetylation status of these ectopic MTs is important for driving this precocious insertion. As previously reported, mRNA injection of the CAMSAP1-Par6 fusion protein leads to a substantial amount of apically positioned MTs (Figure 1G). Importantly, co-injection with HDAC6 significantly lowered the fluorescent intensity of acetylated MTs but not the overall amount of MTs (Figure 1G-J). In addition, the precocious apical insertion observed with CAMSAP1-Par6 is abrogated by the addition of HDAC6 and insertion rates revert to wild type levels (Figure 1K-L) (Collins et al., 2020a). This suggests that MTs in general are not sufficient to promote intercalation but that those MTs must be acetylated.

To further test the effect of acetylated MTs in promoting apical insertion, we OE alpha-tubulin N-acetyltransferase (Tub-ATAT1) to increase MT acetylation in intercalating cells. Tub-ATAT1 expressed specifically in MCCs results in a slight increase in acetylation and a modest but significant increase in cells that apically insert at stage ST18 with this trend continuing at ST20 and ST22 (Figure 2A-F). While the ATAT1 OE result is consistent with our hypothesis that acetylation is important for apical insertion, the already high levels of acetylation in MCCs likely limits the effectiveness. Consistent with this we see a stronger effect on the apical insertion of ICs when we OE ATAT1 using the Pendrin promoter (Pen-ATAT1) which specifically expresses in ICs (Quigley et al., 2011). Pen-ATAT1 OE in ICs resulted in a significant and robust increase in tubulin acetylation and a stronger, significant increase in cells that intercalate at ST20 and ST22 (Fig 2G-L). These results further implicate MT acetylation in promoting apical insertion. We interpret these results to indicate that cells with increased tubulin acetylation have an increased penetrative capacity during radial intercalation.

**Figure 2.**
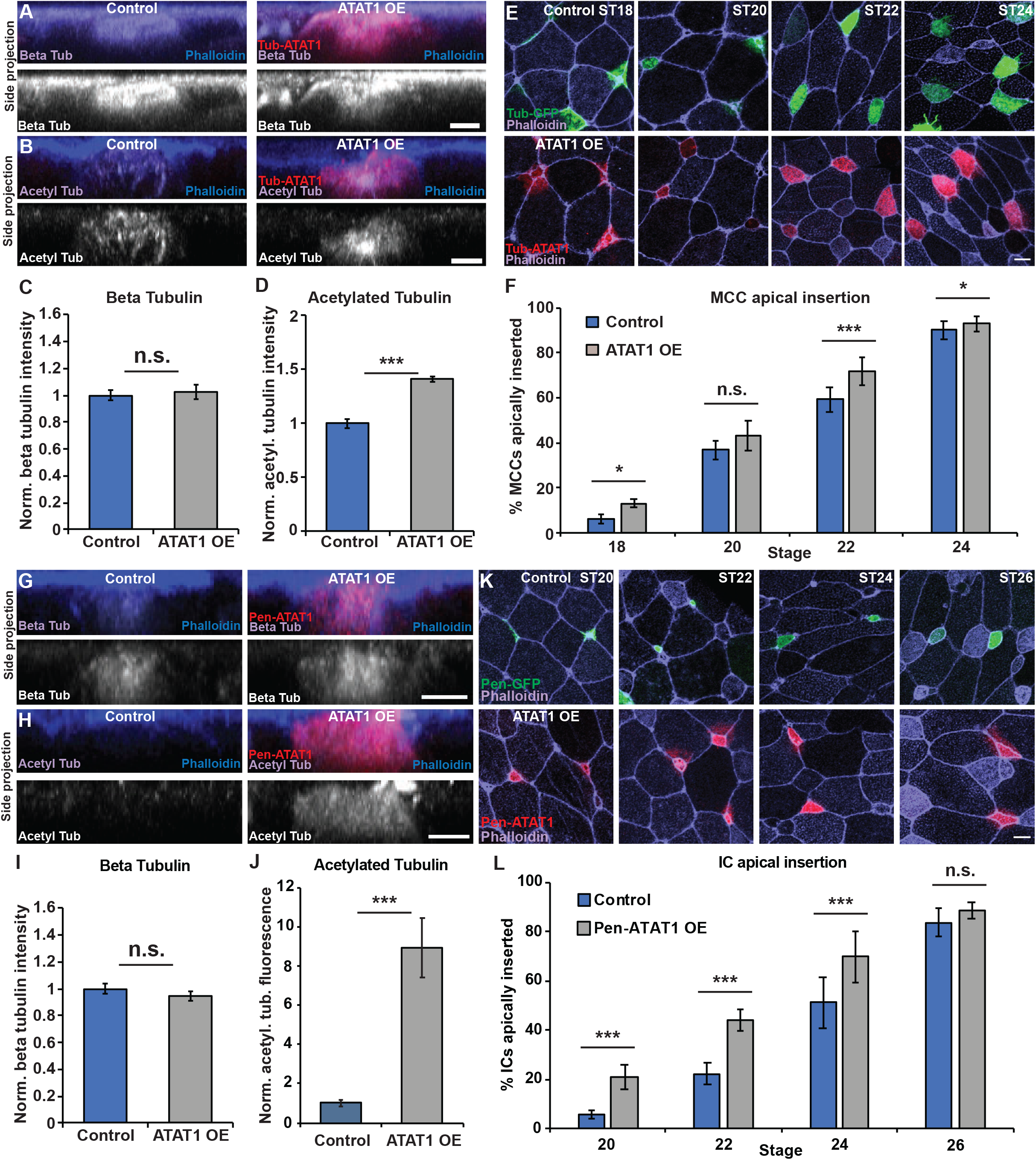
Increased MT acetylation results in precocious apical insertion. **A-B**, Side projections of intercalating control and ATAT1 OE MCCs fixed and stained with α-beta tub **(A)** or α-acetyl. tub. **(B)**. **C**-**D,** Quantification of beta tub **(C)** and of acetyl. tub. **(D)** in control and ATAT1 OE MCCs. Fluorescence was normalized relative to control (uninjected) MCCs in mosaic embryos for each experiment. **E,** Z-projections displaying progression of MCC apical insertion in Control (Tub-RFP) and ATAT1 OE embryos. **F,** Quantification of the percentage of MCCs apically inserted at each stage. **G-H**, Side projections of intercalating control and ATAT1 OE ICs fixed and stained with α-beta tub **(G)** or α-acetyl. tub. **(H)**. **I**-**J,** Quantification of beta tub **(I)** and of acetyl. tub. **(J)** in control and ATAT1 OE ICs. Fluorescence was normalized relative to control (uninjected) ICs in mosaic embryos for each experiment. **K,** Z-projections displaying progression of IC apical insertion in Control (Pen-RFP) and ATAT1 OE embryos. **L,** Quantification of the percentage of ICs apically inserted at each stage. For all bar graphs, bars represent the mean, error bars indicate SD, and *p<0.05 and ***p<0.001. Analysis includes n > 100 cells from at least 6 embryos per condition (C), n > 80cells from at least 5 embryos per condition (D), and n > 200 cells from at least 9 embryos per condition/time point (F), n > 15 cells from at least 3 embryos per condition (I), n > 50 cells from at least 5 embryos per condition (J), and n > 150 cells from at least 7 embryos per condition/time point (L). Scale bars in A,B,G and H is 5μm and in E and L is 10μm.

The process of radial intercalation represents a complex interplay between the intercalating cells and the surrounding outer epithelial tissue. In particular it has been previously reported that cells intercalate exclusively at multicellular vertices rather than between two cells (Stubbs et al., 2006). One interpretation of this observation is that vertices represent weak points in the epithelial barrier that are more easily penetrated. Interestingly, in the complex epithelium of *Xenopus* skin there is a variety of different types of vertices ranging from tricellular junctions to rosettes with 5 or 6 cells (Figure 3A). We hypothesized that different types of vertices would provide differential barrier strength against the penetration of intercalating cells. We performed a live-imaging analysis of vertex preference during the early phases of MCC intercalation (ST17-20). We focused on MCCs specifically because the epithelium is relatively stable during these early stages and MCCs almost never intercalate next to one another which would complicate the interpretation of vertex strength. In contrast, a similar analysis of ICs is not feasible due to the changing vertex environment caused primarily by the intercalation events of the MCCs coupled with the fact that ICs often intercalate adjacent to the sites of MCC intercalation (Drysdale and Elinson, 1992; Dubaissi and Papalopulu, 2011; Stubbs et al., 2006), likely due to the weakened epithelium caused by MCC intercalation.

**Figure 3.**
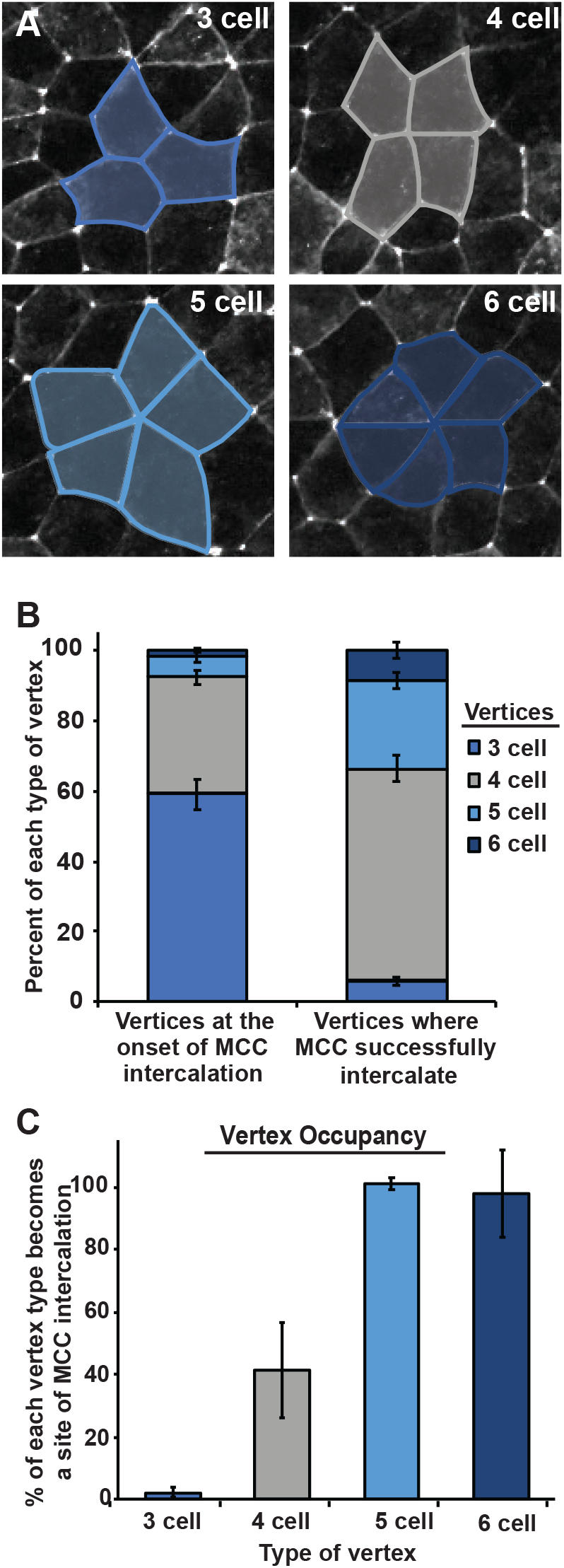
MCCs preferentially intercalate at higher order vertices. **A,** Representative images of 3, 4, 5, and 6 cell vertices between outer epithelial cells expressing GFP-tricellulin (pseduocolored in white). Vertex types are outlined in blue or gray representative of graph colors in **B** and **C**. **B-C,** Analysis of live imaging (e.g. Supplemental Movie 1) quantifying the percentage of each vertex type present in the outer epithelium prior to MCC intercalation (**B**; left) and sites of successful MCC intercalation (**B**; right). **C,** Percentage of each type of vertex occupied with an MCC by stage 20. Analysis includes n > 1500 vertices (**B**, left) and n > 350 MCCs (**B**, right) from 2 embryos, n > 900 vertices (3 cell), 500 vertices (4 cell), 85 vertices (5 cell), and 30 vertices (6 cell) from 2 embryos (**C**).

From our live-imaging movies (e.g. Supplemental Movie 1) we scored the overall number of each type of vertex and found that tricellular vertices were the most common, making up 59% of all vertices (Figure 3A-B). Four cell vertices represented 33% and 5 or 6-cell vertices celled rosettes, accounted for 5% and 2% of vertices respectively (Figure 3B). Interestingly, despite the large number of tricellular vertices we found, MCCs rarely apically inserted at these sites during our live imaging experiments representing only 6% of insertion events (Figure 3B). Four cell vertices, despite being less frequent then tricellular vertices, accounted for close to 60% of all intercalation events (Figure 3B). Finally, while rosettes are rare they account for 34% of intercalation events clearly showing that MCCs have a preference for higher order vertices. In fact, of all 5 and 6 cell vertices, nearly 100% are sites of MCC intercalation between ST17 to ST20 (Figure 3C). In contrast, approximately 42% of all 4 cell vertices and only 3% of all tricellular vertices are sites of successful apical insertion during this period of MCC intercalation (Figure 3C). Interestingly, when the timing of intercalation at each type of vertex is analyzed, a clear pattern arises where cells intercalate earlier at higher order vertices. We scored the cumulative number of cells that intercalate at each type of vertex throughout our time-lapse, and then quantified the time when 50% of those cells had successfully breached the epithelium (T_50_; indicated by the dotted red line in Fig. 4A-D). We found that for 6 cell vertices the T_50_ for MCCs that would eventually occupy a 6 cell vertex was approximately 30 minutes. The T_50_ of 5 cell vertices was slightly slower (~70 minutes), suggesting that these vertices are harder to penetrate. This trend continues with 4 cell vertices having a T_50_ of ~100 minutes and tricellular vertices having a T_50_ of ~160 minutes. The striking regularity of these temporal differences suggest that the challenges of penetrating each type of vertices increased as vertex cell number decreases. Also, our T_50_ quantification suggested a variable strength to the epithelial barrier and we were curious if we could exploit these differences to test the malleability of penetrative capacity for intercalating cells.

**Figure 4.**
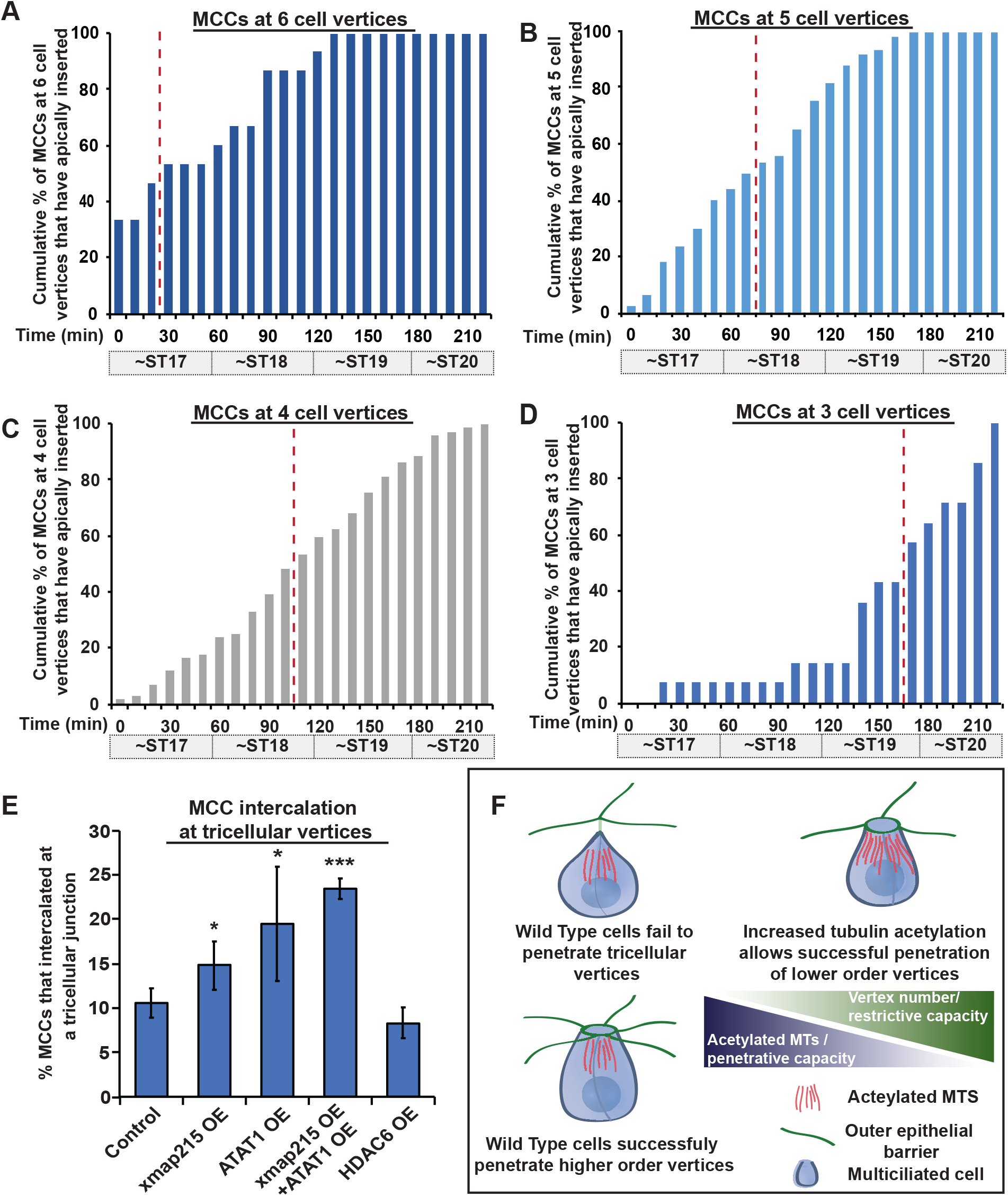
Microtubule acetylation alters penetrative capacity of MCCs. **A-D,** Quantification of cumulative percentages of MCCs inserting in the outer epithelium at 6 cell **(A)**, 5 cell **(B)**, 4 cell **(C)**, and 3 cell vertices **(D)** over time. Percentages were calculated by dividing the cumulative number of total MCCs that had breached the outer epithelial layer at each time point by the number of MCCs that inserted at that vertex type. Red dotted line on each bar graph indicates the point at which 50% of the cells that would eventually intercalate at the vertex type had successfully breached the epithelium. **E,** Quantification of the percentage of MCCs that intercalated at tricellular vertices by stage 20. **F,** Model of the balance of restrictive capacity of the epithelium that varies by vertex strength and the penetrative capacity of intercalating cells that varies by the amount of tubulin acetylation. For bar graph in (E), bars represent the mean, error bars indicate SD, and *p<0.05 and ***p<0.001. A-D are representative data generated from one time-lapse experiment analyzing intercalation of 214 MCCs. Similar trends were seen in other time-lapse videos. Analysis includes n > 350 cells for each condition from at least 9 embryos (E).

Knowing the relative timing of apical insertion for each vertex type gives us a powerful assay to address the relative contribution of tubulin acetylation to the penetrative strength of intercalating MCCs. When analyzing fixed tissues at ST20 we found that only 11% of all MCC intercalation events occurred at tricellular vertices (Figure 4E). We hypothesized that if this low percentage were due to the restrictive strength of the tricellular barrier then this number would be malleable if we altered the penetrative capacity of the intercalating cell by modulating MT number and acetylation. To test this, we first reduced penetrative capacity by injecting embryos with Tub-HDAC6 to lower MT acetylation, and we consistently found that now fewer cells (8%) were able to successfully insert by ST20 (Figure 4E). We next increased MT number by OE the MT nucleating factor XMAP215 and found an increase in both MT numbers and MCCs that successfully intercalated tricellular junctions by ST20 (15%, Figure 4E and Supp Figure 2). Increasing MT acetylation via OE of ATAT1 had an even stronger effect increasing penetration to 20% (Figure 4E and Supp Figure 2). Importantly, OE of both XMap215 and ATAT1 led to an additive increase (23%) of MCCs that were able to apically insert by ST20. These results indicate that increases to MT number and acetylation lead to increased penetrative capacity allowing cells to infiltrate between the restrictive tricellular junctions at a stage (ST20) when the majority of wild type cells fail.

MTs are well known modulators of cell migration and can facilitate the directional movement of membrane vesicles and signaling molecules to the leading edge. Acetylated MTs are more stable and more resistant to strain offering the possibility that they can provide important structural features that deliver strength and rigidity to cells migrating in a 3D tissue environment. Our data provides two important pieces of information that indicate a structural role for MTs in the tissue penetration of radial intercalation. First, in ICs with ectopic apically generated MTs (e.g. CAMSAP1-Par6; Figure 1G-L) the precocious apical insertion is abrogated by the loss of acetylation, indicating that the presence of MTs alone is not sufficient to promote IC apical insertion, but that those MTs must be acetylated. Secondly, the penetration of MCCs at tricellular junctions is promoted separately and additively by the changes to MTs and MT acetylation. These results indicate that during MCC intercalation, not only is the enrichment of MTs important, but that the acetylation of those MTs is also critical. Overall, we propose that MT acetylation facilitates radial intercalation by increasing penetrative capacity by increasing the strength and rigidity of the MT network.

Epithelial tissues provide a barrier function to liquids, toxins, as well as infectious agents. They also provide a barrier to cells attempting to penetrate. Recently, it has been proposed that the topological environment of multiple cells coming together offers easier passage for groups of cells undergoing migration through the drosophila egg chamber (Dai et al., 2020). Our results extend these observations by indicating that this preference for migrating between higher order vertices is a wide-spread evolutionarily conserved feature of diverse tissues with distinct modes of tissue penetration. In addition to bulk collective cell movements, individual cells undergoing radial intercalation also prefer to penetrate at the sites of higher order vertices. More importantly, our results in MCCs indicate that this preference is malleable based on the penetrative capacity of the migrating cell. Consistent with our results, ICs intercalate after MCCs, in part due to lower numbers of centrioles and MTs which would result in lower penetrative capacity. Interestingly, in *Xenopus* embryos, it has been reported that OE of the constitutively active Notch intracellular domain (NICD) leads to a dramatic loss in MCC but not IC fate (Stubbs et al., 2006). In the absence of MCCs there is a significant delay in IC intercalation. This result suggests that the ICs have a weaker penetrative capacity and that they rely on MCC intercalation to create more higher order vertices to facilitate their intercalation. Consistent with this ICs typically intercalate adjacent to MCCs (Stubbs et al., 2006).

There is an ongoing balance between the restrictive competence of the epithelium and the invasive capacity of penetrating cells (Figure 4F). This balance is complex, employing the regulation of junctional remodeling and diverse cytoskeletal elements. Here we provide evidence that one cytoskeletal feature, namely the acetylation of MTs specifically in intercalating cells increases the ability of cells to penetrate by both increasing the overall rate of apical insertion and by facilitating the penetration at more restrictive lower order tricellular vertices.

## Acknowledgments

This work was supported by NIH/NIGMS to BJM (R01GM119322). RV was supported from the Cutaneous Biology training grant (T32AR060710). We would like to thank the National *Xenopus* Resource and the Marine Biological Laboratories for technical support and reagents.

## Experimental Models and Subject Details

*Xenopus laevis* were used and maintained in accordance with standards established by the Northwestern University Institutional Animal Care and Use Committee. Mature *X. laevis* frogs were obtained from NASCO (Fort Atkinson, WI). Frogs were housed in a recirculating tank system with regularly monitored temperature and water quality (pH, conductivity, and nitrate/nitrite levels) at a temperature of 16-18°C and were fed frog brittle. For live imaging and nocodazole experiments, embryos from a Tub-Deup1-GFP transgenic line (Xla.Tg(tuba1a:deup1-eGFP)^NXR^) generated and purchased from National Xenopus Resource RRID:SCR_013731 at the Marine Biological Laboratory were used to identify MCCs.

## Method Details

### Embryo injections

All *Xenopus* experiments were performed using previously described techniques (Werner and Mitchell, 2013). In brief, *Xenopus* embryos were obtained by *in vitro* fertilization using standard protocols (Sive et al., 2007) approved by the Northwestern University Institutional Animal Care and Use Committee. Embryos were injected at the two- or four-cell stage with 40–250 pg mRNA or 10–20 pg of plasmid DNA. For all DNA injections (except for fixed vertices analysis, Figs 3 and 4), embryos were injected mosaically at the 2 or 4 cell stage such that only half of the embryo expressed the construct to avoid toxicity. For fixed MCC vertices analysis (Figs 3 and 4), embryos were injected 3 times in each blastomere at the 2 cell stage for maximal expression of the construct in as many MCCs as possible.

### Plasmids/mRNA

pCS2+ plasmids containing an N-terminal GFP or RFP as a tracer containing the α-tubulin promoter (TUBA1A-B on Scaffold 127187, pCS2tub) (Stubbs et al., 2006) were used to drive expression of some constructs specifically in MCCs. pCS2+ plasmids containing the Pendrin (Pen) promoter (Quigley et al., 2011) with an N-terminal GFP or RFP were used to drive expression of some constructs specifically in ionocytes (Collins et al., 2020a). The GFP-tricellulin construct was described previously, and the GFP-CAMSAP1-Par6 construct has been previously reported (Collins et al., 2020a). HDAC6-FLAG (#13823) and pEF5B-FRT-GFP-aTAT1 (#27099) were purchased from Addgene. Tub-GFP-HDAC6 was made by PCR amplifying the HDAC6 sequence from the HDAC6-FLAG construct and ligating the PCR amplicon into the Tub-GFP construct. MCC- and IC-specific ATAT1 constructs were made by PCR amplifying the ATAT1 sequences and ligating the PCR amplicons into Tub-RFP or Pen-RFP constructs, respectively. An XMAP215-WT-7His-GFP construct was a kind gift from Jay Gatlin and was previously described (Milunovic-Jevtic et al., 2018; Reber et al., 2013). Tub-XMAP215-GFP was made by PCR amplifying the xmap215 sequence and ligating the PCR amplicon into the Tub-XLT vector. mRNA was generated using the Sp6 *in vitro* mRNA transcription kit (Ambion) following linearization of plasmid DNA with NotI. Capped mRNA was isolated using an RNA isolation kit (Qiagen).

### Inhibitors

Nocodazole (Sigma, #M1404) was used to inhibit MT dynamics. Nocodazole treatments were performed between stages 13 and 28. In brief, embryos were incubated in the presence of 0.2 μM or 1 μM Nocodazole (in DMSO) or pure DMSO (vehicle) from ST13 until embryos were at the desired stage. Embryos were fixed in 3% PFA immediately thereafter.

### Immunofluorescence

For antibody staining, embryos were fixed with 3% PFA in PBS, blocked in 10% goat serum, and primary and secondary antibody solutions were prepared in 5% goat serum. Mouse anti-acetylated α-tubulin (T7451; Sigma-Aldrich) was used at a 1:500 dilution, mouse anti-beta tubulin (DHSB; E7) supernatant was used at a 1:10 dilution. Mouse anti-FLAG M2 (F3165; Sigma) and rabbit anti-DYKDDDDK (2368; Cell Signaling) were used at 1:100 dilutions. E7 (anti-tubulin) was deposited to the DSHB by Klymkowsky, M (Chu and Klymkowsky, 1989).. Cy-2–, Cy-3–, or Cy-5–conjugated goat anti-mouse secondary antibodies were used at the manufacturers’ recommended dilution. Phalloidin 650 (1:600, Invitrogen) and Alexa Fluor Plus 405 Phalloidin (1:100, Invitrogen) were used to visualize actin. Embryos were mounted between two coverslips using Fluoro-Gel (Electron Microscopy Sciences).

### Microscopy

All microscopy was performed on a laser-scanning confocal microscope (A1R; Nikon) using a 20× water objective lens (live imaging) or a 60× oil Plan-Apochromat objective lens with a 1.4 NA (fixed imaging). Nikon Elements Software was used for all acquisition and image processing. For all fixed images, multiple z planes were visualized in 0.4 μm steps (4-10 μm total depth). For live imaging, a 20 μm range was imaged with 0.5 μm steps. Images were acquired every 10 minutes for the duration of the time lapse. Images are maximum intensity projections of z stacks. Images were processed in Nikon Elements Software.

## Quantification and Statistical Analysis

### Tubulin intensity analysis

For acetylated and beta tubulin intensity measurements, a z-projection of the intercalating cell was created and the fluorescence intensity was measured in ImageJ by outlining the cell of interest and measuring anti-acetylated or anti-beta tubulin fluorescence within the outline. Tubulin intensities were normalized relative to the mean fluorescence intensity measured in ‘control’ (uninjected cells) in mosaic embryos for each experiment.

### Apical insertion analysis

Apical area of intercalating cells was measured at each stage (based on phalloidin staining). For apical insertion analysis, an apical domain area of 35μm^2^ was set as a threshold to determine apical insertion based analysis previously described (Collins et al., 2020a). The percentage of MCCs or ICs apically inserted (as opposed to still below the surface) at each stage represent the percentage of cells measured at the indicated stage that have an area >35μm^2^ and is independent of measurements taken at other stages. For all apical insertion analyses, cells below the surface of the outer epithelium were included in the analysis and were given an apical area of 0μm^2^.

### Live imaging and MCC vertices analysis

For live imaging analysis, the composition of vertices prior to intercalation was determined by counting the total number of each type of vertex present in the first frame of the time lapse (Fig. 4). To determine the percentages of where MCCs intercalate (Figs. 3 and 4), all MCCs that breached the outer epithelium were analyzed for the type of vertex where they inserted. Vertex occupancy was calculated for each type of vertex by dividing the number of vertices that were a site of MCC intercalation by the end of the time lapse by the total number of each type of vertex counted at the beginning of the time lapse. Bar graphs showing cumulative progression of MCC intercalation at 3, 4, 5, and 6 cell vertices were generated by counting the total number of MCCs that intercalated at each vertex type during the entire time lapse. Cumulative progression of MCC intercalation was calculated at each time point by dividing the cumulative number of MCCs that had inserted at the vertex of interest by the total number of all MCCs that intercalated at that vertex type by the end of the time lapse. The red dotted line on the bar graph represents the time at which 50% of all MCCs that would intercalate at the vertex type had already breached the outer epithelium. Bar graphs displaying cumulative progression of MCC intercalation at different vertex types are representative of data collected for 214 MCCs in one experiment. Similar trends in timing of intercalation 3, 4, 5, and 6 cell vertices were observed in other time lapse experiments. For fixed vertices analyses, embryos fixed at stage 20 were analyzed for sites of MCC intercalation by counting the number of all MCCs expressing the construct of interest that had established an apical domain. The percentage of cells that had intercalated as tricellular vertices was calculated by dividing the number of MCCs expressing each construct that had intercalated at triceullular vertices by the total number of MCCs expressing the construct of interest that had established an apical domain.

### Image presentation and statistical analysis

Some images were smoothed and processed for figure presentation only. Raw images were used for all analyses. Significance was determined with a student’s t-test (Figures 1C, 1D, 1I, 1J, 2C, 2D, 2I, 2J, 4E, S2D) and Chi-square test for (Figures 1F, 1L, 2F, 2L, S1C, S1D). For all bar graphs, bars represent the mean and error bars indicate the standard deviation (SD). For all statistical analyses, *p<0.05, **p<0.01, and ***p<0.001.

## Supplemental Figure Legends

**Figure S1.**
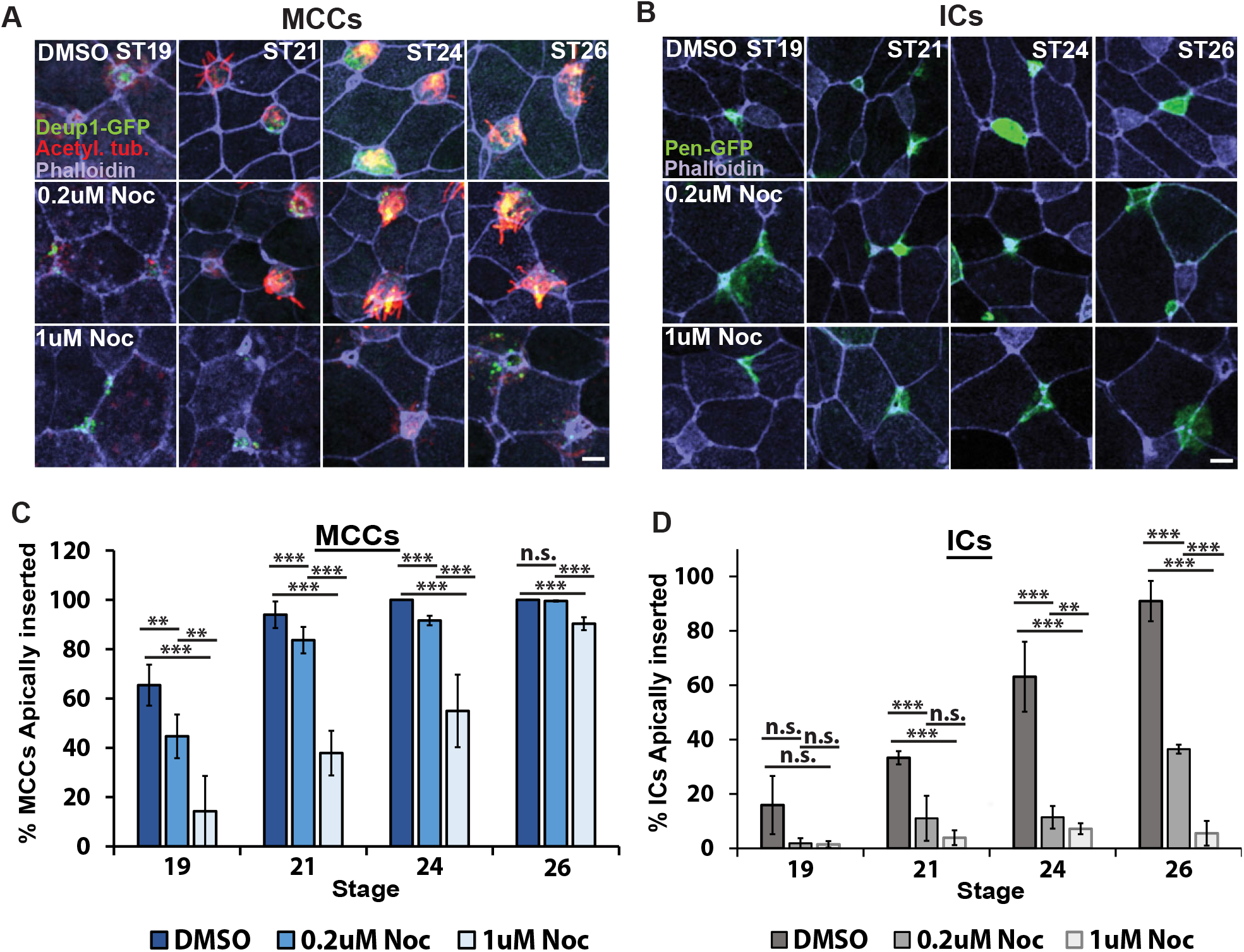
Nocodazole treatment delays MCCs and IC apical insertion in a dose-dependent manner. **A**, Z-projections of Tub-Deup1-GFP embryos treated with DMSO, 0.2μM Nocodazole (Noc), or 1μM Noc fixed and stained with an α-acetylated tubulin antibody and phalloidin at the indicated stages. **B**, Z-projections of WT embryos injected with Pen-GFP and treated with DMSO, 0.2μM Noc, or 1μM Noc fixed and stained with phalloidin at the indicated stages. **C-D,** Quantification of the percentage of MCCs (**C**) or ICs (**D**) apically inserted throughout the intercalation process. Embryos were treated with DMSO, 0.2μM Noc, or 1μM Noc from ST13 until fixation. For bar graphs, bars represent mean and error bars indicate SD, and *p<0.05 and **p<0.01. Analysis includes n > 150 cells at least 9 embryos per condition/time point from (C) and n > 100 cells from at least 9 embryos per condition/time point (D). Scale bars in A-B, 10μm.oint (F), n > 15 cells from at least 5 embryos per condition (I), n > 25 cells from at least 7 embryos per condition (J), n > 50 cells from at least 9 embryos per condition/time point (L). Scale bars in A, B, G and H is 5μm and in E and K is 10μm.

**Figure S2.**
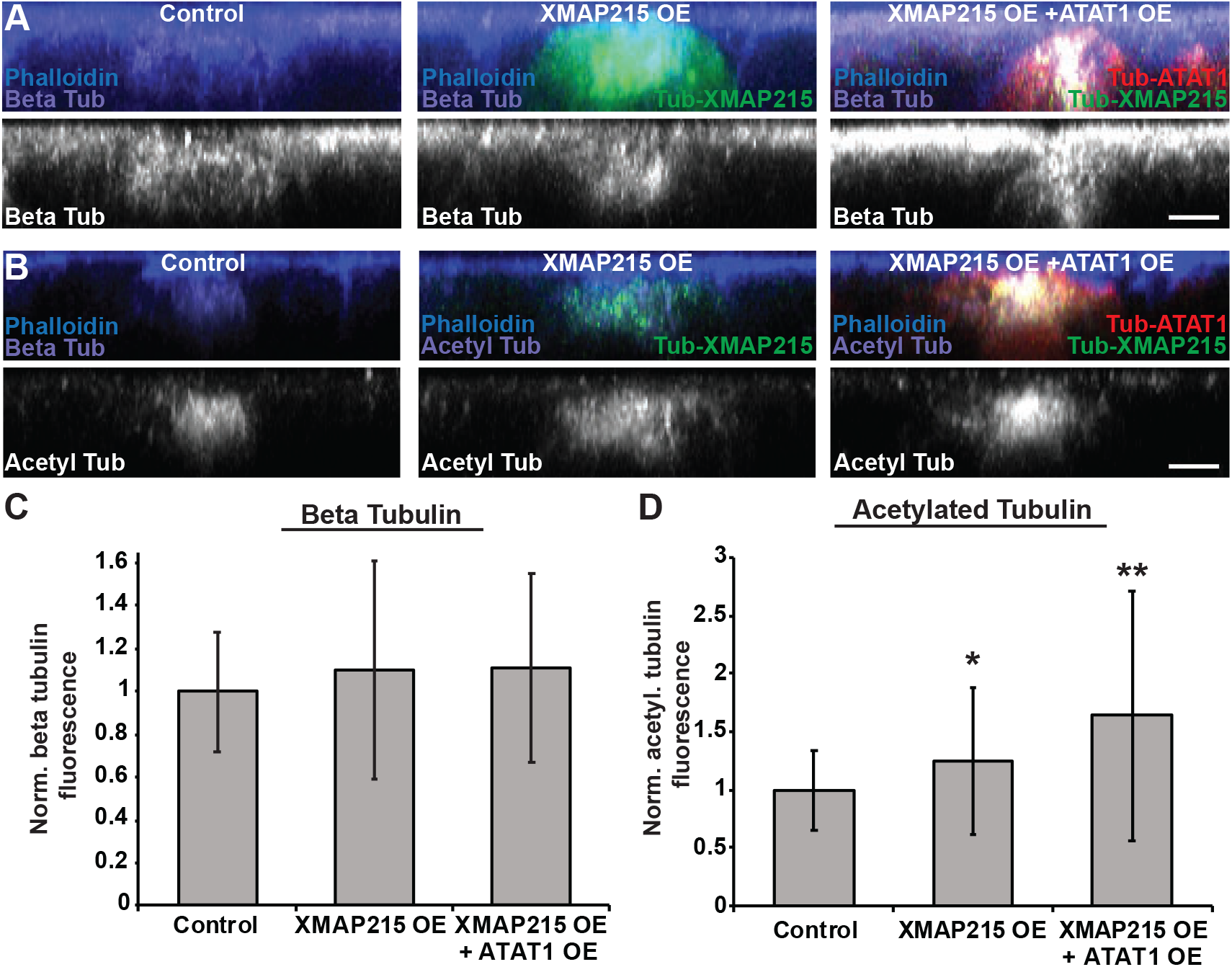
Beta tubulin and acetylated tubulin levels in MCCs expressing MT nucleating and acetylation constructs. **A-B**, Side projections of intercalating control, XMAP215 OE, and XMAP215 OE + ATAT1 OE MCCs fixed and stained with α-beta tub **(A)** or α-acetyl. tub. **(B)**. **C**-**D,** Quantification of beta tub **(C)** and of acetyl. tub. **(D)** in control, XMAP215, and XMAP215 OE + ATAT1 OE MCCs. Fluorescence was normalized relative to control (uninjected) MCCs in mosaic embryos for each experiment. For all bar graphs, bars represent the average, error bars indicate SD, and *p<0.05, **p<0.01. Analysis includes n > 50 cells from at least 6 embryos per condition (C), and n > 45 cells from at least 6 embryos per condition. Scale bars in A-B, 10μm.

---

**Movie S1. Live imaging of MCC intercalation into the outer epithelium.** Time lapse imaging of a Tub-Deup1-GFP transgenic embryo expressing RFP-tricellulin (pseuduocolored white). Deup1-GFP signal was used to identify MCCs. Time lapse imaging began at approximately ST17. A 20 μm z-range was imaged (0.5 μm steps) every 10 min throughout the duration of the time lapse.

